# The response of genes and pathways of immunological system induced by irradiation

**DOI:** 10.1101/526384

**Authors:** F. M. Ribeiro, D. A. Silveira, E. M. Simão, V. L. D. Mattos, E. G. Góes

## Abstract

Current studies have shown that ionizing radiation (IR) could increase the efficiency of radiation therapy by the stimulation of the immune system. This occurs in low-dose radiation as well as doses within hypofractionated range usually used in radiotherapy. However, the elucidation of the mechanisms of immunogenic modulation reported at these doses remain an issue. In this study, we analyzed transcriptome data available in Gene Expression Omnibus (GEO) database related to B cells isolated from whole blood of 95 donors and then irradiated with 10 Gy. The aim of this study is to investigate the regulation of genes and pathways of the immune system considering the B7-CD28/CTLA4 superfamily, CD40-CD40LG molecules, and cytokines expressed by B cells irradiated. The connection between genes and pathways is established by the Reactome database. Relative activity and diversity of pathways were calculated to determine the modulation of the immune system response to irradiation. Analysis of variance (ANOVA) with repeated measures and Bonferroni’s method were used to determine differentially expressed genes. It was observed that IR up-modulates the response of pathways and genes considered in this study, which indicates that 10 Gy can enhance antitumor immune responses.

## Introduction

The earliest applications of ionizing radiation (IR) in medicine were accomplished immediately after Roentgen discovery of x-rays in 1895. Since then, new technologies and methodologies associated with the use of IR in healthcare have been developed to maximize the benefits from its use in detriment of undesirable effects. In medicine, these applications focus on diagnostic radiology, nuclear medicine, and radiation oncology. Researches of radiation oncology biological basis have shown that cell response to IR can be described in terms of the damage produced by radiation in DNA^1^. In this way, the radiobiological model proposed by Withers consider that the damage induces different biological responses related to damage repair, redistribution in the cell cycle, accelerate repopulation, reoxygenation, and intrinsic radiosensitivity, denoted as the “5 R’s of radiobiology”^1–4^. Such effects represent the basis of radiobiology relates to optimization of the therapeutic process used to treat cancer using IR^5^.

Besides the direct damage to cancer cells, there is also clinical evidence that RIs inhibit tumor growth outside treatment field^5–9^. After irradiation, most cells survive a limited period; however, this time is long enough to generate molecular signals necessary to trigger events associated with the immunological response^10–12^. In this sense, our group decided to investigate effects of the dose of 10 Gy on pathways and genes associated with molecules of B7-CD28/CTLA4 superfamily. In addition, considering the role of cytokines in the radiation-induced immune response is not well understood, the effects of this dose on the modulation of the response of genes related to cytokines expressed by B cells were also investigated by us.

## Methods

### Microarray data selection

We selected transcriptome data related to B cells isolated from whole blood of 95 unrelated donors and irradiated with 10 Gy. These data are available in the Gene Expression Omnibus (GEO) database and designated by the access number GSE36910. In order to analyze microarray data, 285 samples were collected and arranged in 3 groups, each containing 95 samples, according to the time considered after irradiation: zero (control), 2h, and 6 h. The design of the experiment and a detailed description of the methods used, as well as the details about the dataset, can be found in the original publication^13^ Affymetrix website was used to get the information related to the HG-U133_2 array (platform GPL571) to determine the gene that is associated with each probe set.

### Robust Multi-array Average Method

Transcriptome data was processed by the Robust Multi-array Average (RMA) method using the R software (https://www.r-project.org/). This process was performed in 3 steps: corrections related to background noise, normalization, and summarization of the probe response associated with the level magnitude of the gene expression^14,15^. The ath1121501 package with Chip Description File (.CDF) was used to identify the position of each probe on the microarray surface. In this context, the Bioconductor affy package was used to access and process the information associated with microarray data stored on CEL files. The highest expression value was used in those cases where multiple probes represented the same gene, according to Stalteri & Harrison^16^ recommendation. All these steps were performed through Bioconductor packages, available in http://bioconductor.org/.

### Relative activity and diversity calculation

In this study, it was considered 44 immune system pathways containing genes associated with molecules of the B7-CD28/CTLA4 superfamily (Table 1) available from Reactome database^17,18^. We used the model described by Castro and coworkers to calculate the relative activity and relative diversity of the pathway as a function of the dose of 10 Gy^19^. According to this model, relative activity (nα) is used to measure an increase or decrease in genes cumulative activity in a pathway α, in relation to control:

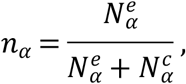

where, 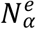 and 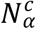 are the activity of the α pathway associated with experiment and control, respectively^20^, in which nα values are restricted between 0 ≤ *nα* ≤ 1. For a group of genes, relative diversity was used to compare data distribution associated with the pathway in relation to the control, through the following relation:

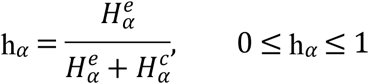

where, 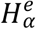 and 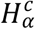 are the Shannon entropy of the *α* pathway associated to the experiment and control, respectively. The Shannon’s entropy *H_α_*, also called diversity, is defined through the following equation:

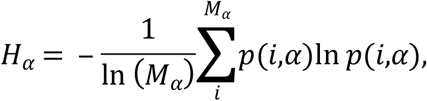

where, ln (*M_α_*) is the normalization factor used to guarantee that 0 ≤ *H_α_* ≤ 1^19,21^.

**Table 1.** Response of the immune system pathways in function of 10 Gy after 2 and 6 h of exposure: increase (p<0.05) in relative activity values (nα) or relative diversity values (hα) is indicated by upward arrows (↑), decrease (p≥0.95) of these values is indicated by downward arrows (↓), and (—) indicates that the pathway was not altered by irradiation (p>0.05).

| Immune system pathways (Reactome database) | Number of genes | Genes associated to superfamily observed in each pathway | n<sub>α</sub> |  | h<sub>α</sub> |  |
| --- | --- | --- | --- | --- | --- | --- |
|  |  |  | 2 h | 6 h | 2 h | 6 h |
| CD28 co-stimulation | 32 | CD28, CD80, CD86 | — | — | — | — |
| CD28 dependent PI3K/AKT signaling | 75 | CD28, CD80, CD86 | — | — | — | — |
| CD28 dependent VAV1 pathway | 12 | CD28, CD80, CD86 | — | — | — | — |
| Costimulation by the CD28 family | 75 | CD28, CD80, CD86, CTLA4, ICOS, ICOSLG, PDCD1, PDCD1LG2 | — | — | — | — |
| CTLA4 inhibitory signaling | 22 | CD80, CD86, CTLA4 | — | — | — | — |
| DAP12 interactions | 340 | CD28, CD80, CD86 | — | — | — | — |
| DAP12 signaling | 325 | CD28, CD80, CD86 | — | — | — | — |
| Downstream signaling events of B cell receptor (BCR) | 150 | CD28, CD80, CD86 | ↑ | ↑ | — | — |
| Downstream signaling of activated FGFR1 | 310 | CD28, CD80, CD86 | — | — | — | — |
| Downstream signaling of activated FGFR2 | 310 | CD28, CD80, CD86 | — | — | — | — |
| Downstream signaling of activated FGFR3 | 310 | CD28, CD80, CD86 | — | — | — | — |
| Downstream signaling of activated FGFR4 | 310 | CD28, CD80, CD86 | — | — | — | — |
| FC epsilon receptor (FCERI) signaling | 388 | CD28, CD80, CD86 | — | — | — | — |
| GAB1 signalosome | 86 | CD28, CD80, CD86 | ↓ | — | — | — |
| Immune system | 1129 | CD28, CD80, CD86, CTLA4, ICOS, ICOSLG, PDCD1, PDCD1LG2 | — | ↓ | — | — |
| Immunoregulatory interactions between a lymphoid and a non lymphoid cell | 94 | CD80 | — | ↓ | — | ↑ |
| Innate immune system | 644 | CD28, CD80, CD86 | ↑ | — | — | — |
| NEF mediated downregulation of CD28 cell surface expression | 3 | CD28 | — | — | — | — |
| NGF signaling via TRKA from plasma membrane | 355 | CD28, CD80, CD86 | — | — | — | — |
| PDCD1 signaling | 21 | CD28, CD80, CD86 | — | ↑ | — | — |
| PI3K ATK activation | 85 | CD28, CD80, CD86 | ↑ | ↑ | — | ↓ |
| PI3K cascade: FGFR1 | 332 | PDCD1, PDCD1LG2 | ↑ | ↑ | ↓ | ↓ |
| PI3K cascade: FGFR2 | 100 | CD28, CD80, CD86 | — | — | — | — |
| PI3K cascade: FGFR3 | 100 | CD28, CD80, CD86 | — | — | — | — |
| PI3K cascade: FGFR4 | 99 | CD28, CD80, CD86 | — | — | — | — |
| PI3K events in ERBB2 signaling | 83 | CD28, CD80, CD86 | ↑ | ↑ | — | ↓ |
| PI3K events in ERBB4 signaling | 83 | CD28, CD80, CD86 | ↑ | ↑ | — | ↓ |
| PIP3 activates ATK signaling | 83 | CD28, CD80, CD86 | ↑ | ↑ | — | ↓ |
| Role of LAT2 NTAL LAB on calcium mobilization | 99 | CD28, CD80, CD86 | ↑ | ↑ | — | — |
| Signal transduction | 1665 | CD28, CD80, CD86 | — | ↑ | — | — |
| Signaling by EGFR | 335 | CD80, CD86 | — | — | — | — |
| Signaling by ERBB2 | 320 | CD28, CD80, CD86 | — | — | — | — |
| Signaling by ERBB4 | 311 | CD28, CD80, CD86 | — | — | — | — |
| Signaling by FGFR | 317 | CD28, CD80, CD86 | — | — | — | — |
| Signaling by FGFR1 | 314 | CD28, CD80, CD86 | — | — | — | — |
| Signaling by FGFR2 | 316 | CD28, CD80, CD86 | — | — | — | — |
| Signaling by FGFR3 | 313 | CD28, CD80, CD86 | — | — | — | — |
| Signaling by FGFR4 | 313 | CD28, CD80, CD86 | — | — | — | — |
| Signaling by NGF | 432 | CD28, CD80, CD86 | — | — | — | — |
| Signaling by PDGF | 345 | CD28, CD80, CD86 | — | — | — | — |
| Signaling by SCF-KIT | 306 | CD28, CD80, CD86 | — | — | — | — |
| Signaling pathway | 180 | CD28, CD80, CD86 | ↑ | — | — | — |
| TCR signaling in NAIVE CD4 T cell | 44 | CD28, CD80, CD86 | — | ↑ | — | — |
| TCR signaling in NAIVE CD8 T cell | 43 | CD86, PDCD1 | — | ↑ | — | — |

The Via Complex software was used to perform relative activity and relative diversity calculations, and statistical calculation as well. The Bootstrap method^22^ was used in random resampling with 10,000 repetitions for all the genes of each transcriptome and pathway to determine whether if the alteration in relative activity or relative diversity was statistically significant. The data were reported as mean ± 1SD (n=95), and a significant level of α = 0.05 was considered for all the statistical analysis. In this sense, it was considered that the irradiation produced a significant increase in relative activity or in relative diversity for p≤0.05. Also, it was considered that irradiation produced a significant decrease in relative activity or in relative diversity for p≥0.95. For values in the range of 0.05<p<0.95, it was considered that irradiation did not change the relative activity or the relative diversity of the pathway.

### Interaction networks, and gene fold change analysis

Gene interaction networks^23^ were used to analyze the genes systematic behavior in response to irradiation. These interaction networks were built using the String database (https://string-db.org/), considering the following interactions: databases, text mining, and experiments. The level of reliability related to the interactions was established at 0.7, according to the literature^24^. Fold change value was used to determine individually the response of the gene to irradiation, represented in the network by the Cytoscape software - version 3.2.1 (http://www.cytoscape.org/). For each gene, the fold change is calculated based on the ratio of expression between experimental e control groups^25^.

Differences in expression levels between irradiated and control groups were tested by repeated measures ANOVA. The null hypothesis H0 (sphericity) was tested using the Mauchly sphericity test. The Greenhouse-Geisser method was used in those cases of rejection of the H0 hypothesis (H1 alternative hypothesis)^26^. In cases where it was observed significant changes in expression levels caused by irradiation, the Bonferroni method was used to make sure that these changes did not happen by chance. All statistical analysis of data were accomplished using the IBM SPSS Statistics 22 software^27^. The data were reported as mean ± 1SD (n=95), and a significance level of α = 0.05 was considered by us in all statistical analysis.

## Results

### Pathways alterations

According to the data obtained from the Reactome database, 44 cellular pathways related to the immune system were selected, which contained genes associated with molecules of the B7-CD28/CTLA4 superfamily (Table 1). It was observed that 10 Gy could up regulated or down regulated the expression of 16 pathways (Table1). According to these data, irradiation produced a significant increase (p≤0.05) in the relative activity, as well as a decrease in the relative diversity of the pathways: PI3K cascade, FGFR1, PI3K events in ERBB2 signaling, PI3k events in ERBB4 signaling, PI3K/ATK activation, and PIP3 activates ATK signaling. However, 10 Gy also produced an increase in the relative activity (p≤0.05) of the pathways: downstream signaling events of B cell receptor, innate immune system, signal transduction, TCR signaling in naive CD4, TCR signaling in naive CD8, LAT2 NTAL LAB function on calcium mobilization, signaling pathway, and PDCD1 signaling. On the other hand, 10 Gy produced a significant decrease (p≥0.95) in the relative activity of pathways: immune system, GAB1 signalosome, and immunoregulatory interactions between lymphoid and non-lymphoid. Among these pathways, just immunoregulatory interactions between lymphoid and non-lymphoid presented a significant increase in the relative diversity (p ≤ 0.05). Despite these effects, this dose did not modify the response of 28 pathways, as shown in Table 1.

### Response of molecules of B7-CD28 / CTLA4 superfamily to irradiation

Table 2 presents levels of genes expression associated with molecules of the B7-CD28/CTLA4 superfamily as a function of exposure time (2 h or 6 h) to 10 Gy. Levels of genes expression are reported in Table 2 using mean of ± 1.0 SD. According to these results, it was observed that 10 Gy produced an increased in the expression level of the ICOSLG gene in both 2 and 6 h after exposition (p ≤ 0.05). However, the expression level of the CD28 gene increased 2 h after irradiation (p = 0.011) but not in 6 h (p = 0,453). On the other hand, the expression level of the CD86 gene increased only 6 h after irradiation (p = 0.017). The expression level of the CD80 gene remained unchanged at 2 h after irradiation (p = 0.026) but showed a significant increase at 6 h after irradiation (p = 0.028). The PDCD1LG2 and PDCD1 genes were the only ones which presented a decrease in the expression levels after 2 or 6 h of irradiation (p≥0.95). According to our results, the expression level of the PDCD1 gene remained unchanged at 2 h after irradiation but showed a decrease (p = 0.001) at 6 h after irradiation. On the other hand, PDCD1LG2 gene presented a decrease in expression level (p = 0.047) 2 h after irradiation but remained unchanged 6 h after irradiation. It was also observed by us that 10 Gy did not change the expression levels of the CTLA4, ICOS, and VTCN1 genes (p> 0.05). In addition to the superfamily of molecules, we also decided to determine the genes expression response to 10 Gy related to CD40, and CDLG molecules. It was observed by us an increase in the expression levels (p≤0.05) of these genes at 2 and 6 after irradiation.

**Table 2.** Effects of 10 Gy on genes expression levels related to B7-CD28/CTLA4 superfamily, CD40, and CD40LG after 2 and 6 h of irradiation: increased expression (p≤0.05) is indicated by the upward arrows (↑), decreased expression (p≥0.95) is indicated by downward arrows (↓), and NA indicates that 10 Gy did not change the expression of the genes (p>0.05) when compared to control (zero hour).

| Genes | Probe set | Expression levels after irradiation |  |  | Status of the expression after irradiation |  |
| --- | --- | --- | --- | --- | --- | --- |
|  |  | (mean ± 1SD; n = 95) |  |  | 2 hours | 6 hours |
|  |  | Control (zero hour) | 2 hours | 6 hours |  |  |
| CD28 | 206545_at | 6.14 ± 0.97 | 6.26 ± 0.99 | 6.09 ± 0.93 | ↑ | NA |
| CD40 | 205153_s_at | 9.85 ± 0.32 | 9.99 ± 0.30 | 9.86 ± 0.35 | ↑ | NA |
| CD40LG | 207892_at | 6.08 ± 0.23 | 6.09 ± 0.27 | 6.11 ± 0.28 | NA | NA |
| CD80 | 207176_s_at | 9.03 ± 0.33 | 8.87 ± 0.31 | 9.28 ± 0.33 | NA | ↑ |
| CD86 | 210895_s_at | 9.21 ± 0.81 | 9.23 ± 0.75 | 9.30 ± 0.71 | NA | ↑ |
| CTLA4 | 221331_x_at | 5.56 ± 0.20 | 5.59 ± 0.20 | 5.55 ± 0.18 | NA | NA |
| ICOS | 210439_at | 4.26 ± 0.24 | 4.28 ± 0.24 | 4.26 ± 0.24 | NA | NA |
| ICOSLG | 213450_s_at | 7.66 ± 0.28 | 7.87 ± 0.32 | 7.84 ± 0.27 | ↑ | ↑ |
| PDCD | 207634_at | 5.64 ± 0.22 | 5.65 ± 0.24 | 5.58 ± 0.21 | NA | ↓ |
| PDCD1LG2 | 220049_s_at | 6.09 ± 0.42 | 5.93 ± 0.35 | 6.14 ± 0.47 | ↓ | NA |
| VTCN1 | 219768_at | 4.98 ± 0.17 | 4.97 ± 0.15 | 5.01 ± 0.19 | NA | NA |

### Genes Expression Response associated to cytokines related to B cell

In this study, we also considered the interaction network modeling of 27 cytokines related to B cell (Figure 1). In this modeling, the size of the molecule is proportional to the fold change value of each gene that makes up the network. As state in this network, 10 Gy of irradiation produced an increase in the expression levels (p≤0.05) of the following genes: IL10RA, IL4R, IL10RB, IL1R1, TGFB1, TGFBR1, TGFBR2, IFNGR2, IFNG and IL1A (Figure 1, and Table 3). On the other hand, this does also produce a significant decrease (p≥0.95) in the expression levels of the following genes: IL10, IL6R, IL2RB, IL2RG, IFNGR1, IL12RB2, IL1R2, IL12A, and IL12B (Figure 1, and Table 3).

**Figure 1.**
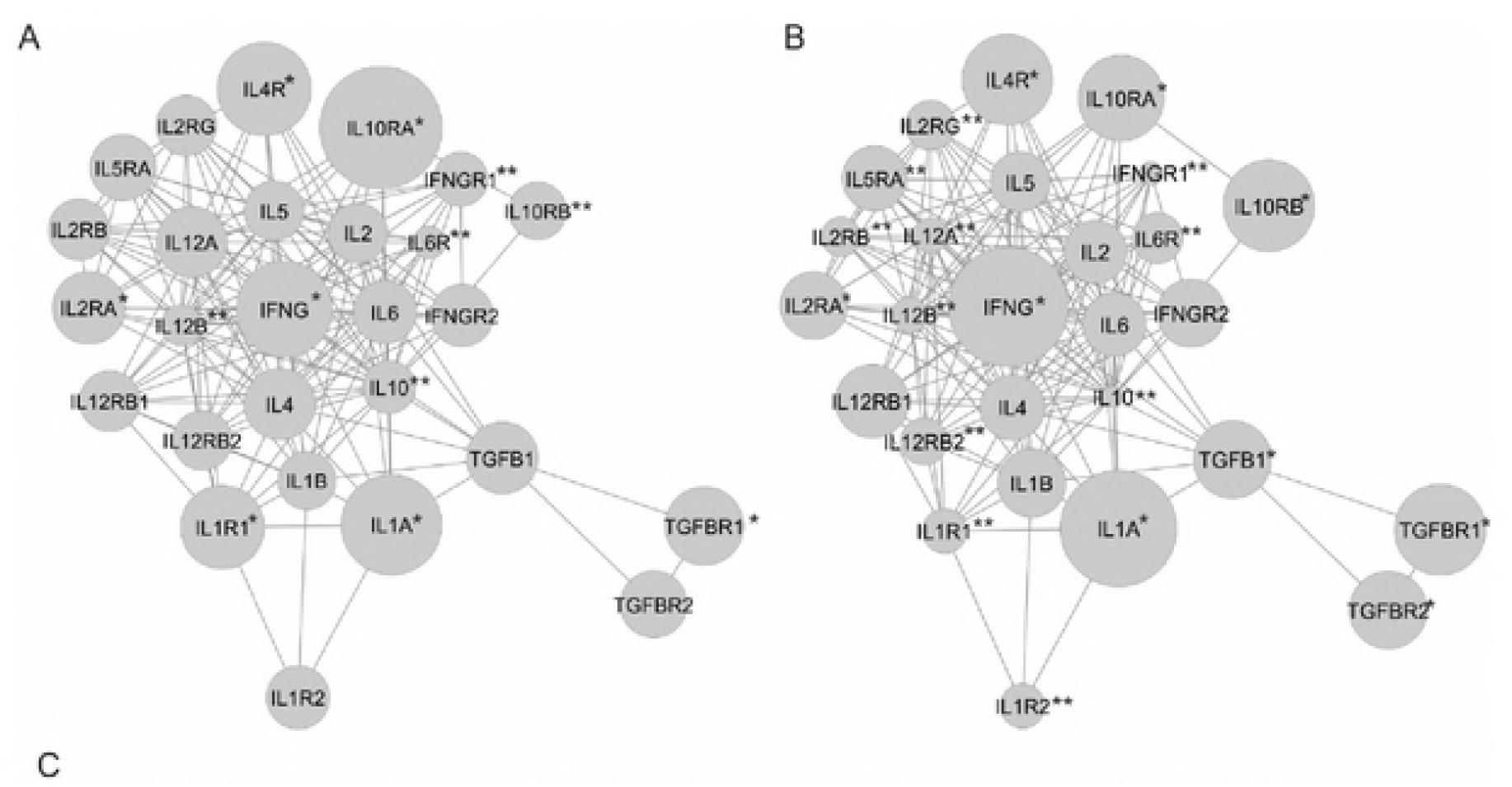
Interaction network of 27 cytokines related to B cells. The size of each node represents the magnitude of the irradiation effect on gene expression measured (mean ± 1SD; n = 95) after 2 hours (A) and 6 hours (B) of irradiation. The values of expression of each gene are shown in Table 3. The symbol * indicates that 10 Gy increase the expression of the gene (p≥0.95), and the symbol ** indicates that 10 Gy decrease the expression (p > 0.05), at 2 or 6 hours after irradiation when compared to the control (zero hour).

**Table 3.**
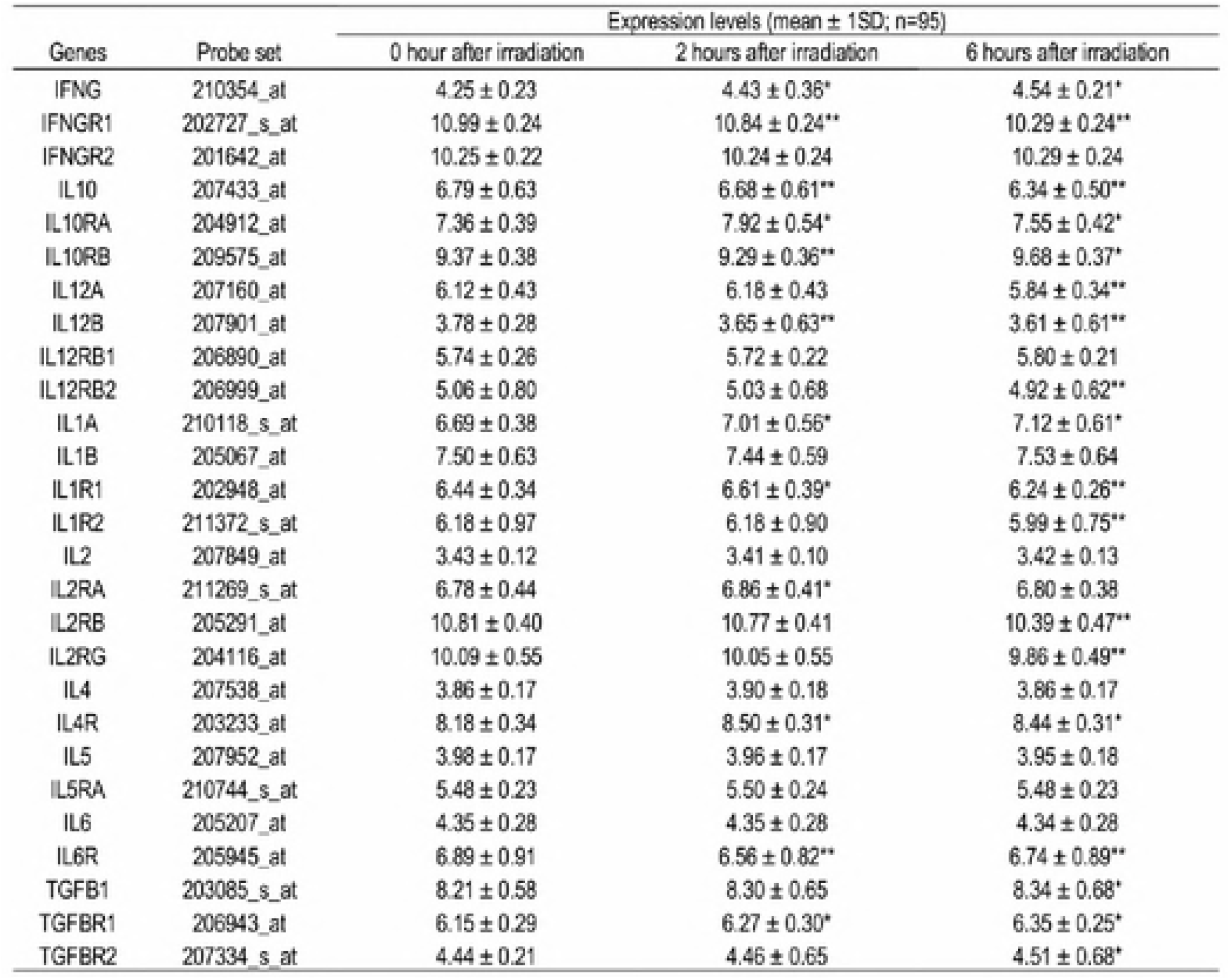
Effects of 10 Gy on the levels of gene expression related to cytokines expressed in ß cells The symbol * indicates that 10 Gy increased the levels of genes expression (p≤0.05), and the symbo ** indicates that 10 Gy decreased this expression (p≥0.95), at 2 or 6 hours after irradiation wher compared to the control (2er0 hour).

## Discussion

The results obtained in this study provide an overview of the regulation of immune response by irradiation. 44 pathways of the immune system were investigated which containing genes related to molecules of the superfamily, 16 were observed to be significantly modified by irradiation regarding the relative activity (Table 1). Based on these results, 13 pathways had a cumulative activity of the genes, while a decrease in this activity was observed in 3 pathways. Thus, the irradiation up-regulated and down-regulated 29,5% and 6,8% of the pathways investigated, respectively. In addition, according to results related to the relative diversity calculation (Table 1), the irradiation promoted genes differentially expressed in 5 pathways (11,4%); one pathway up-regulated, and 4 pathways down-regulated. These results indicate that 10 Gy modulated in a differentiated way the organization of the biological system. An important concept of system reorganization is the up-regulation of the TCR signaling pathways in naïve CD4 and CD8 T cells (Table 1), and the MCH II antigen pathway associated with B cell. These findings are important because these pathways are able to promote the activation, proliferation, and differentiation of cells related to the immune system. As reported by Topalian, these events are necessary for the induction and sustention of the immune response^28^. In agreement with two-signal model, are required two signals for the activation of the immune response: initialization, the first signal, and maintenance (and enlargement) of the response, the second signal^29–31^. The first signal is associated with T cell receptor recognition and T cell binding to MHC antigen, which is presented by APCs. The second signal is associated with the occurrence related to both cellular and molecular events, including T cell signaling, differentiation, and proliferation^32,33^. In this study, B cells respond to irradiation according to its biological mechanism that resembles this model. In this sense, the up-regulation of MHCII antigen pathway activates the first signal, and the up-regulation of the TCR signaling pathways in naïve CD4 and CD8 T cells activates the second signal. In addition, the results showed that 10 Gy enriched genes differentially expressed (up or down-regulated) related to cytokines (Figure 1), and to the molecules of the B7-CD28/CTLA4 superfamily (Table 2). These conclusions are associated with the second signal, necessary to activate the immunological response, in agreement with the two-signal model previously described.

It is necessary both stimulatory/costimulatory and inhibitory/coinhibitory second signals to modulate TCR-mediated T cell receptor activation^34^. In this study, as presented in Table 2, 10 Gy produced a significant increase in the expression level of the CD28 gene. This molecule provides costimulatory signals which are essential for amplification and maintenance of T block response through interaction with its CD80 and CD86 ligands, which are expressed on cell surface, as in B cells^11^. Moreover, 10 Gy also produced a significant increase in the expression levels of the CD80 and CD86 genes (Table 2). As reported by the literature, the interaction of the B7-CD28 molecules emits a positive signal that, together with TCR signaling, promotes the activation, proliferation, and differentiation of T cells^35^. In this context, the results obtained in this study indicate that irradiation induces interaction of the B7-CD28 molecules, activating T cells, which are essential for the initiation and maintenance of humoral and cellular immune response mediated by specific antigens. In addition, it was observed by us that 10 Gy produced a significant increase in the expression level of the ICOSLG gene, which is expressed in the surface of B cells (Table 2). This molecule expresses its immunological functions binding to its ICOS receptor^36^. The ICOS/ICOSLG interaction contributes to T cells costimulatory signal which results in T cells amplification and B cells activation, increase of proliferation, and cytokine secretion^37^. Studies associated with the functions of this surface molecule have demonstrated that the ICOS signaling plays an essential role in the differentiation of Th1 and Th2 cells by activation of the specific antigens, as well as in the promotion of Th17 response^37,38^. Another strong costimulatory signal is provided by the CD40 molecule, which performs its immunological functions through the interaction with its CD40LG ligand. As shown in the results, 10 Gy produced a significant increase in expression level of the CD40 gene. The CD40 is a molecule expressed in APCs which plays an important function on control of B cells response with dependence in T cells. In this context, the interaction of B and T cells could promote B cell proliferation, survival, immunoglobulin production, and B cell memory formation^39,40^. The interaction between CD40 molecule and the CD40LG ligand regulates an important molecular and cellular processes associated to the adaptive humoral immunity^41,42^. Based on these results, this study indicates that irradiation induce an increase in expression level related to the CD40 molecule in the B cells surface, which stimulates the activation and differentiation of these cells. Furthermore, 10 Gy could induce the amplification of the B7-CD28 interaction, necessary for the expansion and maturation of effectors T cells^43,44^.

According to the results presented in Table 2, 10 Gy did not produce significant changes in the expression level of the CTLA4 gene. The CTLA4 surface molecule is a CD28 analogue, with substantial B7 (CD80 and CD86) binding affinity. However, unlike CD28, this negative regulatory molecule could inhibit T cell responses^45^. This inhibitory signal could produce mechanisms, such as TCR signaling inhibition, CD28 molecule inhibition, and/or inhibition of it signaling pathway. Consequently, the B7-CTLA4 complex decreases the ability to interact between T cells and APCs, such as B cells^46^. The CTLA4 molecule, unlike CD28, is expressed at low levels in resting T cells and resides mainly in the cytoplasmic matrix47. This molecule is expressed on the surface of the activated T cells and has the role of regulating down the immune response. CTLA4 ability to inhibit T cell activation depends on the intensity of TCR signal, and B cell activation. In this context, the present study does not exclude that 10 Gy could induce B7-CTLA4 interaction; thus, regulating negatively T cell activation, which decreases T cell proliferation. The CTLA4 molecule executes this function on up-regulate T cells surface, not on B cells, which explain the fact that no change in CTLA4 gene expression level was observed by us. Similar to the CTLA4 molecule, PDCD1 is an inhibitory molecule that plays an important role in the regulation of the immune response. Recent studies have shown that PDCD1 is an immune control protein from T cell which negatively regulates the immune response^48^. The PDCD1 receptor has three ligands: PDCDLG1 (Programmed Death Ligand-1), PDCD1LG2 (Programmed Death Ligand-2), and CD80. The PDCD1 limits the activity of T cells by inducing a phosphatase that inhibits the receptor-mediated signal^49^. In this study, 10 Gy irradiation produced significant decrease on PDCD1, and PDCD1LG2 genes expression levels (Table 2). As stated in the literature, CTLA4 and PDCD1 are negative regulators with non-redundant function for the modulation of immune response^50^. T cells, B cells, and monocytes activation induce the expression of PDCD1 gene^51^. The interactions between PDCD1 and its ligands could inhibit the proliferation of T cells, the INFG, the tumor necrosis factor, the IL2 production, and reduces T cell survival^46^. In this context, the results of this study indicate that the 10 Gy inhibits the PDCD1/PDCD1LG2 interaction signal, thus up-regulating the immune response.

As reported by Muller and collaborators, the production of cytokines is related to the third sign necessary to activate the effector phase of the immune response^52^. Cytokines represent a diverse group of small soluble proteins that could function as factors of growth and differentiation in the autocrine or paracrine forms. Through binding to specific cell surface receptors, they initiate signal transduction pathways that are crucial for a varied spectrum of functions, including induction of immune responses, cell proliferation, and differentiation, for example. In this study, we analyzed the expression of 27 genes related to cytokines expressed in B cells in response to irradiation (Figure 1). It was observed by us that the irradiation significantly modified the expression of 19 genes associated with these cytokines (p≤0.05). Based on these results, 10 Gy produced a significant increase in the expression level of IL1R1 gene. On the other hand, this dose of irradiation produced a significant decrease in the expression level of IL1R2 gene. The IL1R1 receptor-mediated signaling stimulates the activation of IL1 cytokine, which is necessary for the initiation and maintenance of the immune system activities, such as B and T cells differentiation, for example^53^. When expressed on the surface of B cells, the second receptor (IL1R2) blocks the IL1 transduction signal which inhibits the IL1 cytokine function^54,55^. The IL1R1 receptor activation, and the IL1R2 receptor inhibition on B cells surface could stimulate its maturation, differentiation, and co-stimulation, as well as an increase in antibodies synthesis^56^. Based on the results presented in figure 1 and table 3, 10 Gy also produced a significant increase in expression level of IL2RA gene and a significant decrease in expression levels of IL2RB and IL2RG genes, which are related to their respective receptors located on the cell surface of both activated B and T cells^57^. The production of IL2 cytokine could be stimulated by cells that express the dimer and trimer of mean (IL2RB and IL2RG) and high affinity (IL2RA, IL2RB and IL2RG), respectively^58^. IL2 signs could trigger immune response, presenting differentiation and homeostasis of lymphocytes. Maintenance of regulatory T cells (Treg), and CD4 T cells differentiation depends on the signal related to IL2. In addition, signals related to IL2 optimize T cells production, and CD8 differentiation in T cells memory^58^. As stated on the literature, activation of IL2R receptors on B cells surface promotes B cell proliferation, which is stimulated with CD40/CD40LG interaction^59^. Furthermore, to increase B cells differentiation the IL2/IL2R interaction should cooperate with other cytokines/stimulatory-factors^60,61^. The IL4 and the IL6 are other cytokines that could activate B cell differentiation. Based on the results, 10 Gy irradiation produces a significant increase in IL4R gene expression level, which is the receptor expressed on the B cells surface associated with IL4 cytokine (Figure 1). The receptor activation on the B cells surface increases the proliferation induced by the CD40/CD40L interaction, and receptor binding. Also, IL4/IL4R interaction induces a class shift of immunoglobulin, preferably for IgG1, IgG4, and IgE^62,63^. On the other hand, as shown in the results presented in Figure 1, 10 Gy of irradiation produced a significant decrease in the expression level of the IL6R gene. This gene is associated with the IL6 receptor, expressed on the surface of the B cell. Inhibition of this receptor on the B cells surface decreases the differentiation of these cells^64^. It was also observed by us that 10 Gy produced a significant decrease in the expression level of the IL10 gene, which is expressed on the cellular surface of activated B cells (Figure 1). This indicates an activation of the immune system. IL10 is one of the most important cytokines related to the anti-inflammatory process, it could inhibit proinflammatory responses of innate and adaptive immune cells^65^. Cytokines production in macrophages could increase with the inhibition of these cytokine, growth and suppression of lymphocyte-mediated immunity could occur with the inhibition^64^. On the other hand, 10 Gy of irradiation produced a significant increase in receptors expression levels associated with the IL10, IL10RA, and IL10RB, expressed on the B cells surface (Figure 1). Moreover, the activation of these receptors causes proliferation, B cells differentiation, and changing of immunoglobulin class, associated with IgG1 and IgG3^66–68^. The IL10 interaction with receptors expressed on B cells surface collaborates with the TGFB cytokine to induce a class change in association with IgA. The signal contributes to B cells differentiation in plasma, which secretes IgM, IgG and IgA^64,68^. Based on the results presented in Figure1, 10 Gy of irradiation produced a significant decrease in the expression level of IL12 gene. The production of this cytokine induces an increase, differentiation, promotion of cytotoxic CD8 T cell. Also, it induces activation, proliferation, and NK cells production of IFNG^64^. Furthermore, signal associated with IL12 contributes to B cell differentiation in IgM-secreting cells. On the other hand, this signal suppresses IgE production induced by IL4^68^. Based on the results, 10 Gy of irradiation produced a significant increase in the expression levels of the IFNGR2 gene. This dose significantly decrease the expression levels of the IFNGR1 gene, which express receptors on the B cells surface associated with IFNG (Figure 1). The activation of these receptors promote on the B cell surface a class change of IgG2a immunoglobulin ^64,69^. It was also observed that 10 Gy produced a significant increase in the levels of expression of the TGFBR1 and TGFBR2 genes, which express receptors on the B cells surface associated with the TGFB1 ligand (Figure 1). Activation of these receptors on the B cells surface promotes IgA synthesis (CD79A and CD79B)^64^.

The main contribution of cytokines to B cell differentiation is its ability to modulate the expression of these transcription factors, which regulates Ig secretion using activated B cells^64^. In this context, the costimulatory action produced by the B7-CD28 interaction enhances the previously activated B and T cell responses, promoting the production of interleukin-2 (IL2) and T cell survival^11^. After initiation and differentiation of lymphocytes, the production of effectors cytokines, such as IL4, does not require costimulatory action of the B7-CD28 interaction. The IL2 production depends on the costimulatory signaling^10,44^. In this context, B7-CTLA4 interaction reduces IL2 production, and IL2 receptor (IL2R) expression. Recent studies have shown that ionizing radiation induces the activation of pathways with pro- and anti-proliferative signal, which imbalance the decision of the cellular destination^70^. Radiation is able to regulate the expression of genes and factors involved in cell cycle progression, cell survival or death, DNA repair and inflammation modulating a radiation-dependent intracellular response^71^. In radiotherapy, irradiation modulates anti-tumor immune responses by modifying of the tumor and its microenvironment^72^. The balance of pro-inflammatory and anti-inflammatory cytokines is important to determine the positive or negative result, adverse reactions, and treatment resistance^73^. Radiation therapy has a significant effect on the modulation of the immune system through the activation of cytokine cascades^74^. Cytokine signature analysis associated with B cell is, therefore, a study of interest for the understanding of the role of cytokines in the radiation-induced immune response. The results obtained in the present study provide an overview of the expression of genes related to interleukins, represented on the surface of the microarray HG-U133_2, in function of the dose of 10 Gy released in B cells.

## Conclusions

Relative activity and diversity is a mathematical model that has been proposed to describe the behavior of pathways in cancerous tissues through transcriptome data. In this type of study, biological networks, their modeling, visualization, and analysis have been used to describe biological models related to cancer. This methodological approach was used by us in order to study the response of the immune system to irradiation. The results observed by us indicated that 10 Gy up-regulated most of the pathways considered here. Also, it was observed a significant increase in expression levels of genes related to co-stimulatory signals and a significant decrease in the expression of the genes related to the inhibition of costimulatory signals. Both processes are necessary to trigger and maintaining the immune response. Furthermore, an increase in expression levels of the genes related to cytokines that induce lymphocyte activation, differentiation and proliferation has also been observed in this study. All these findings indicate that 10 Gy up-regulates the immune system response.

## Acknowledgments

This research was partially supported by CAPES. This work is based in a MSc thesis part submitted for the partial fulfillment of the requirements of the graduate programme in physics at the Federal University of Rio Grande, Rio Grande, Rio Grande do Sul, Brazil.

